# Unconventional Translation Initiation Factor EIF2A is required for Drosophila spermatogenesis

**DOI:** 10.1101/2021.03.28.437419

**Authors:** David D. Lowe, Denise J. Montell

## Abstract

The eukaryotic initiation factor EIF2A is an unconventional translation factor required for initiation of protein synthesis from non-AUG codons from a variety of transcripts, including oncogenes and stress related genes in mammalian cells. Its function in multicellular organisms has not been reported. Here, we identify and characterize mutant alleles of the CG7414 gene, which encodes the Drosophila EIF2A ortholog. We identified that CG7414 undergoes sex-specific splicing that regulates its male-specific expression. We characterized a Mi{Mic} transposon insertion that disrupts the coding regions of all predicted isoforms and is a genetic null allele, and a PBac transposon insertion into an intron, which is a hypomorph. The Mi{Mic} allele is homozygous lethal, while the viable progeny from the hypomorphic PiggyBac allele are male sterile and female fertile. In dEIF2A mutant flies, sperm failed to individualize due to defects in F-actin cones and failure to form and maintain cystic bulges, ultimately leading to sterility. These results demonstrate that EIF2A is essential in a multicellular organism, both for normal development and spermatogenesis, and provide an entree into the elucidation of the role of EIF2A and unconventional translation i*n vivo*.

## Introduction

Protein synthesis, or mRNA translation, is a complex and fundamental biological process. A key point of regulation is initiation because it acts as the rate-limiting step. Therefore, translational initiation plays a significant role in regulating the proteomic diversity of a cell. Canonical translation initiation in eukaryotes begins with an AUG start codon, or translation initiation site, which is recognized by the ribosome scanning across the 5’ UTR^1^ (Fig. 1A). Recently, bioinformatic and modified mass spectrometry analysis^2–3^ have identified unconventional or non-canonical translation initiation at alternative (i.e. non-AUG) start codons. Accumulating evidence suggests that translation initiation at non-AUG start codons is a mechanism of translational regulation in eukaryotes, especially during stress, and its mis-regulation is common in disease^4^.

**Figure. 1.**
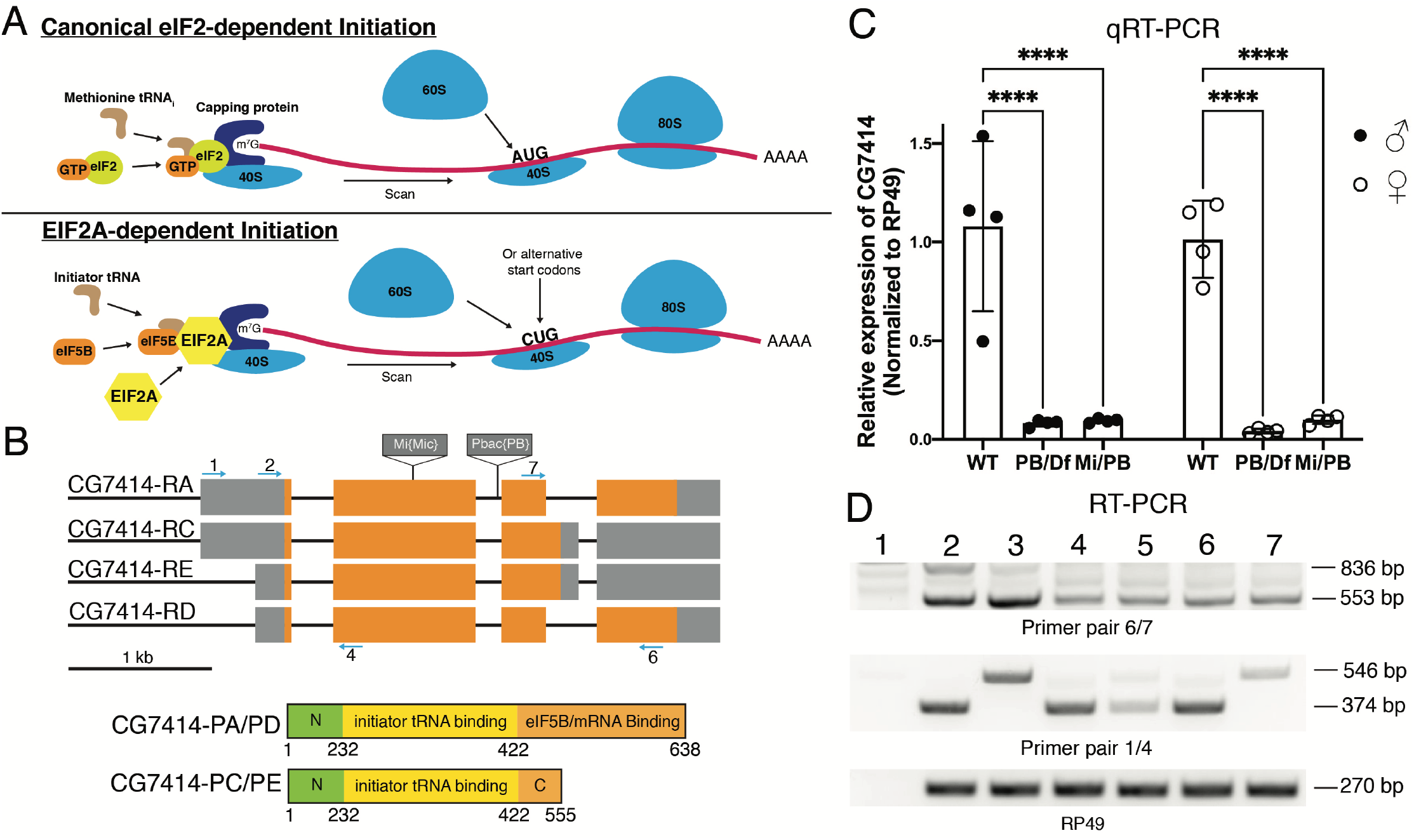
EIF2A schematic and analysis of CG7414 mutant alleles. (A) Schematic representation of eIF2 vs. EIF2A-dependent translation. (B) CG7414 transcripts are shown with sites of transposon insertion. Coding sequence in orange and UTRs in gray. Below, the two protein isoforms are shown with differing C-terminus. Location of the primers used for PCR are shown as cyan arrows. (C) qRT-PCR is shown with primer pair 1/2. CG7414 expression was normalized to RP49. Relative expression of CG7414 was normalized to Wild-type or WT of the respective sex. Df represents *Df(3L)BSC284*. Four biological replicates for each sample. **** p-value <0.0001 (D) RT-PCR is shown. Lane 1: RT(-) as negative control; Lane 2: WT Male; Lane 3: WT Female; Lane 4: PB/*Df(3L)BSC284* Male; Lane 5: PB/*Df(3L)BSC284* Female; Lane 6: Mi/PB Male; Lane 7: Mi/PB Female. Primer pair 6/7 leads to 846 bp band representing CG7414-RC/RE, while the smaller 553 bp band represents CG7414-RA/RD transcripts. Primer pair 1/4 leads to 374 bp band that represents the spliced transcripts, while the larger 546 bp band represents the un-spliced CG7414-RA/RC transcripts expressed in female flies. RP49 serves as loading control.

Eukaryotic initiation factor 2A (EIF2A) is a specialized translation initiation factor for initiation at CUG or UUG start codons instead of the canonical AUG^5^ (Fig. 1A). Analysis of EIF2A in mammalian systems has shown its importance for tumorigenesis and synthesis of CUG initiated peptides^6^. In mice, the PTEN tumor suppressor has a functional CUG-initiated isoform that is EIF2A-dependent^7^. Additionally, the long isoform of the p38 activated protein kinase MK2 is CUG initiated^8^. In Drosophila, the protein dFMR1 has a functional CUG initiated isoform that is expressed in the nervous and reproductive systems^9^. The initiation of translation at CUG codons is evolutionarily conserved from flies to humans.

Canonical translation initiation is primarily regulated by the GTP-dependent eIF2 complex, which is composed of 5 subunits. The eIF2 complex binds GTP and the methionine initiator tRNA to form the eIF2 ternary complex, which then binds the 40S ribosome to form the 43S pre-initiation complex^10^. The target mRNA transcript to be translated contains a m^7^G cap at the 5’ end and poly A tail at the 3’ end that work synergistically for translation^11,12^. The capping protein recognizes the m^7^G cap and recruits the 43S pre-initiation complex for cap-dependent translation initiation. Once bound to the transcript, the pre-initiation complex scans along the mRNA transcript until it finds a functional AUG start codon to recruit the 60S ribosome^13,14^. In addition, EIF5B promotes GTP hydrolysis to dissociate the eIF2 complex to progress towards translation elongation and 80S ribosome formation^15^.

On the other hand, EIF2A-dependent translations have different requirements that allow for translation initiation at non-AUG start codons. EIF2A can function independent of GTP to bind the initiator tRNA and form an alternative ternary complex compared to the GTP-dependent eIF2 complex. The alternative ternary complex with EIF2A then binds with the 40S ribosome to form the 43S pre-initiation complex that scans along the 5’UTR until it finds a functional non-AUG start codon (Fig. 1A). In the case of EIF2A, CUG and UUG are viable non-AUG start codons^5^. In addition, eIF5B binds to the C-terminus of EIF2A likely to dissociate the GTP-independent EIF2A to allow for translation elongation similar to its role in canonical translation initiation^16^ (Fig. 1B). However, the detailed mechanism of EIF2A-mediated translation is not well understood due to the lack of structural, biochemical, and genetic data available compared to the eIF2 complex.

EIF2A is highly conserved across eukaryotes, however the role of EIF2A *in vivo* in any multicellular organism has yet to be reported. Here, we report that the Drosophila ortholog of the EIF2A gene is CG7414. We show that null alleles in CG7414 are predominantly lethal, whereas hypomorphs are 100% male sterile. We report male-specific splicing of the first intron. Furthermore, males lacking sufficient expression of CG7414 failed to produce mature spermatids because of the absence of waste bags and the dysfunctional cystic bulges due to abnormal individualization complexes. This study demonstrates that dEIF2A is essential for viability and male fertility and opens the door to detailed mechanistic studies *in vivo*.

## Results

### CG7414 is essential for viability and male fertility

*CG7414* is a single copy gene that that is predicted to produce four transcripts, two of which are highly expressed in the testis (RA/RC) and two of which are widely expressed (RD/RE) ^17^. The CG7414-RA and - RD mRNA transcripts encode a 638 amino acid polypeptide considered to be the full-length protein, whereas CG7414-RC and -RE encode a 555 amino acid isoform (Fig. 1B). The two polypeptides differ only at the carboxy termini and are approximately equally related to the human protein (Fig. S1A; 36% identical for CG7414-PA/D and 34% identical for CG7414-PC/E). They both contain the beta-propeller-like domain necessary for initiator tRNA binding activity (Fig. S1A). However, the polypeptide encoded by the CG7414-RA and -RD mRNAs contains the conserved C-terminal mRNA- and predicted eIF5B-binding domain identified in mammalian EIF2A (Fig. S1B), whereas the polypeptide encoded in CG7414-RC and -RE mRNA transcripts does not^16^ (Fig. 1B).

To assess the function of CG7414, we identified and characterized mutations in the locus. The PBac{PB}CG7414^c00723^ allele, hereafter labeled as PB, contains a PiggyBac transposon insertion into the second intron (Fig. 1B). The Mi{MIC}CG7414^MI06462^ allele, hereafter Mi, contains a MiMIC transposon insertion into the second exon (Fig. 1B). Homozygous PB flies were semi-lethal (31/831 viable progeny), while Mi was homozygous lethal (0/361). When crossed with deficiencies *Df(3L)BSC284* or *Df(3L)ED4978*, which span the CG7414 gene, Mi was lethal demonstrating a requirement for dEIF2A for organismal viability in Drosophila (Table 1). PB was viable over both deficiencies, and since CG7414-RA and RC mRNA transcripts are highly expressed in males, we assessed the requirement of CG7414 for fertility. Males carrying the PB allele in trans-heterozygous combination with either of the two deficiency lines were sterile, whereas the females were fertile (Table 1). In addition, the rare PB homozygous mutants were also male sterile and female fertile (Table 1). These results suggest that the PB is a hypomorphic allele while Mi is a genetic null. We conclude that CG7414 is required for viability and male fertility.

**Table 1.** Viability and fertility defects in CG7414 mutants.

| Genotype | Viability | Male fertility | % of Male fertility | Female fertility |
| --- | --- | --- | --- | --- |
| Mi/Mi | Lethal | - | - | - |
| PB/PB | Semi-lethal | Sterile | 0%, N=9 | Fertile |
| Mi/PB | Viable | Sterile | 1%, N=96 | Fertile |
| *PB/Df(3L)ED4978* | Viable | Sterile | 0%, N=37 | Fertile |
| *PB/Df(3L)BSC284* | Viable | Sterile | 0%, N=38 | Fertile |
| *Mi/Df(3L)ED4978* | Lethal | - | - | - |
| *Mi/Df(3L)BSC284* | Lethal | - | - | - |

### Transposon insertions reduce expression of CG7414

To assess the effects of the transposon insertions on CG7414 mRNA expression, we isolated total RNA from wild-type and CG7414 mutant flies. Each sample was separated by sex to determine sex-specific effects on gene expression. The RNAs were reverse transcribed with oligo d(T), and primers were designed to amplify the testis-enriched transcripts, all transcripts or full-length and truncated transcripts (Fig. 1B). To quantify the effects on mRNA expression, we performed RT-qPCR using primer pair 2/4 to target the first exon, which is present in all predicted isoforms. Both PB/*Df(3L)BSC284* and Mi/PB male and female flies exhibited significant decreases in CG7414 mRNA compared to wild-type male and female flies, respectively (Fig. 1C). Thus, the transposon insertions significantly reduced expression of dEIF2A.

To evaluate which CG7414 transcripts were expressed in wild-type and mutant males and females, we use primer pair 6/7 (Fig. 1B). A 553 bp PCR product representing CG7414-RA and -RD and a larger 836 bp product representing CG7414-RC and - RE was amplified from both male and female flies (Fig. 1D). The larger 836 bp PCR product was predominantly expressed in males (Fig. 1D; Lane 2) and were reduced in the mutant lines (Fig. 1D; Lanes 3-7). Altogether, these results suggest that the transposon insertions reduced the expression of CG7414.

### CG7414 undergoes sex-specific splicing

To distinguish the CG7414 isoforms annotated as testis-enriched from those annotated as ubiquitous, we designed primer pair 1/4, which targets the extended 5’ UTR that is annotated to be present only in testis-enriched isoforms -RA and -RC (Fig. 1B). The expected 374 bp PCR product was observed in wild-type males (Fig. 1D; Lane 2). Surprisingly, a larger 546 bp product was amplified from wild-type females (Fig. 1D; Lane 3). Sequencing of the gel-extracted products revealed retention of 172 nucleotides corresponding to the first intron in RNA from female but not male flies (Fig. S2). This is in agreement with previous work reporting sex-specific retention of introns in many genes, including CG7414^18^. In addition, we noticed the presence of faint larger bands in PB/*Df(3L)BSC284* and Mi/PB male flies (Fig. 1D; Lane 4 and 6) and the appearance of the smaller spliced product in female flies (Fig. 1D; Lane 7). Thus the transposon insertions alter splicing as well as overall expression levels of the transcripts^19^. We conclude that CG7414 transcripts undergo sex-specific splicing and thus sex-specific post-transcriptional regulation of protein expression. Altogether, the data show that CG7414 is required for normal development and for male fertility, and that male fertility is more sensitive to reductions in CG7414 expression.

### CG7414 is required for spermatid maturation

To further characterize the male sterile phenotype, we examined the male reproductive system. Spermatogenesis occurs in the testis, which is a reproductive structure in male flies that resembles a coiled tube. Distinct transcriptional and translational phases regulate spermatogenesis at different developmental stages^20,21^. At the apical end of a testis, there is a hub of 6-12 germline stem cells (GSCs; Fig. 2A) which divides asymmetrically into two cells: a daughter stem cell and a differentiating gonialblast (GB). The GB is flanked by two somatic progenitor cyst cells which envelope the GB. The GB goes through four cycles of mitosis with incomplete cytokinesis to form a cyst of 16 primary spermatocytes (Fig. 2A). In the early stage, primary spermatocytes will expand their cell volume up to ~25 fold and initiate transcription of mRNAs for translation during meiosis. In the late stage, primary spermatocytes will proceed through meiosis I and II resulting in the generation of 64 haploid spermatids^22^ (Fig. 2A). The round haploid spermatids processes a mitochondrial derivative termed nebenkern that elongates with the axoneme^23^. During elongation, the spermatids will polarize and undergo changes in nuclear shape and chromatin condensation. The elongated spermatid will remodel through a caspase-dependent individualization process to form individualized mature sperm^24^, which are transported to the seminal vesicle for storage until copulation (Fig. S3A). Interestingly, sperm maturation requires multiple, specific translation initiation factors^25^.

**Figure. 2.**
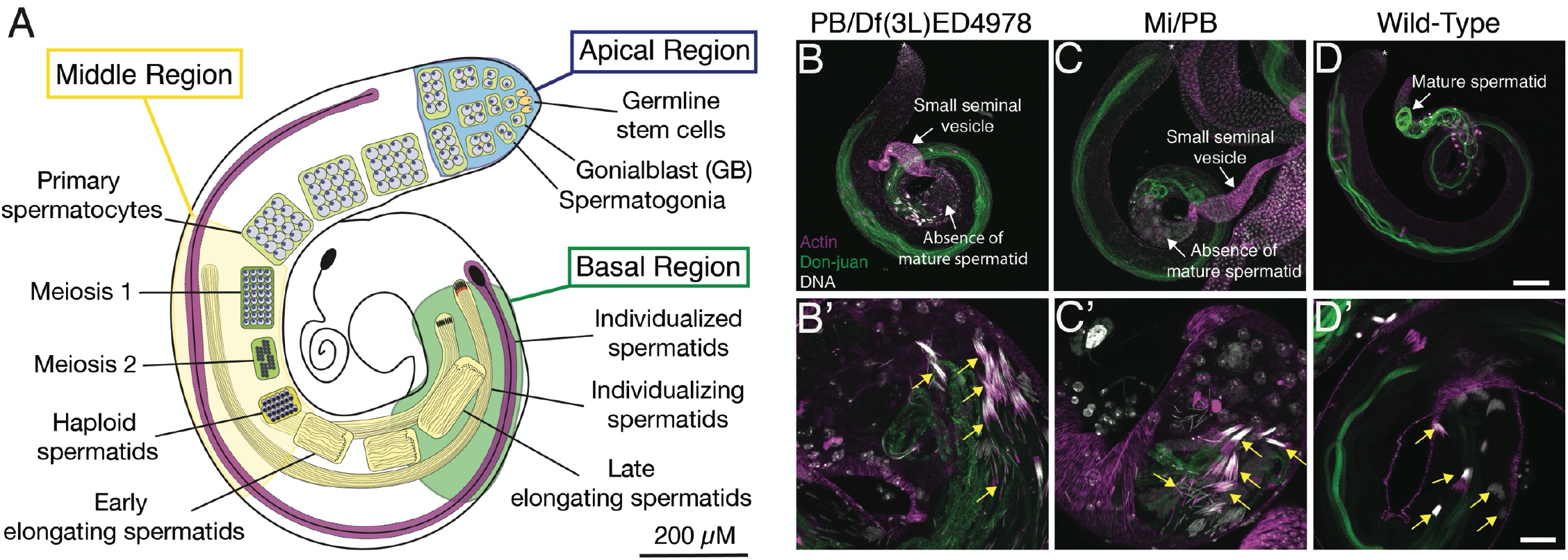
Defective spermatogenesis in CG7414 mutants. (A) Outline of spermatogenesis in testes with black scale bar at 200 μM. (B) PB/*Df(3L)ED4978* testis are shown at 10X magnification and (B’) at 40X magnification with yellow arrows showing assembling individualization complexes with condensed nuclei and actin cones. (C) Mi/PB are shown at 10X magnification and (C’) at 40X magnification with yellow arrows showing assembling individualization complexes with condensed nuclei and actin cones. (D) Wild-type testis are shown at 10X and (D’) at 40X with yellow arrows showing assembling individualization complexes with condensed nuclei and actin cones. (B-D’) Actin staining is labeled as magenta, DJ-GFP is labeled as green, and Hoechst staining is labeled as white. A white asterisk marks the apical end of the testis. Scale bars: 100 μM, and 25 μM.

Light microscopy of homozygous PB/PB testes showed a small seminal vesicle compared to the normally large seminal vesicles in WT and heterozygous PB/+ testes (Fig. S3 A-C). Furthermore, confocal microscopy showed a similar small seminal vesicle phenotype in both Mi/PB and PB/*Df(3L)ED4978* testis, suggesting a reduction or absence in the production of mature sperm in CG7414 mutants (Fig. 2B, C) compared to wild-type (Fig. 2D). Therefore, we investigated what stage(s) of spermatogenesis was (were) disrupted. Spermatogenesis was examined with GFP-tagged Don-juan protein (DJ-GFP), a mitochondrial protein previously characterized to be post-meiotically expressed and a marker for late spermiogenesis^26^. We used DJ-GFP to label elongating spermatids and mature sperm. Both Mi/PB and PB/*Df(3L)ED4978* testis exhibited weak signals of DJ-GFP and an absence of mature spermatid in the basal end of the testis (Fig. 2B-D; white arrows) compared to WT which had strong DJ-GFP signal and mature sperm in the basal end of the testis (Fig. 2D; white arrow). These observations suggest that the male sterility from the loss of CG7414 is due to the absence of mature sperm.

We next determined if elongation was affected by the loss of CG7414 and looked for the presence of needle-like nuclei and F-actin that correspond to late stages of elongation prior to individualization^27^. Both Mi/PB and PB/*Df(3L)ED4978* testis displayed the presence of needle-like nuclei and assembly of F-actin cones (Fig 2B’, C’; yellow arrows) similar to the WT (Fig. 2D’; yellow arrows). These results suggest that elongation proceeds similarly in CG7414 mutants and wild-type and that the loss of CG7414 specifically affects the later stages of spermiogenesis, such as individualization.

### Loss of cystic bulge and waste bags in CG7414 mutants

Individualization is a highly orchestrated process that employs the apoptotic machinery including caspase enzymes to remodel spermatids into mature sperm^28^. Near the end of elongation, the nuclei of 64 immature spermatids condense into needle-like shapes with dense actin structures and assemble a macromolecular structure called the individualization complex. The individualization complex is composed of a coordinated array of F-actin cones or investment cones (Figure 2D’; yellow arrows). As the individualization complex begins to mobilize, the F-actin cones become triangular and polarized and drive the remodeling of the spermatids from the basal end to the apical tip of the testis^27,29^. Once the individualization complex starts to migrate, it becomes a visible, dense bulge termed the cystic bulge, which accumulates organelles, mitochondria, and cytoplasmic components during individualization. At the end of individualization, the accumulated cytoplasmic bulge pinches off from the cyst to form a waste bag to be degraded near the apical tip of the testes.

To look at different stages of individualization complexes, testes were immunostained for both cleaved Death Caspase-1 (cDcp-1) to label active executioner caspases, and phalloidin to label F-actin. As expected, cDcp-1 was present in individualization complexes outlining the individualizing spermatids in the WT testes (Fig. 3A, A’). Initial assembling individualization complexes were characterized as spermatids with co-localized Hoechst and phalloidin staining, which is characteristic of late elongation stages when individualization complexes are near the nuclei (Fig. 2D’; yellow arrows), as shown previously^27,29^. Cystic bulges were characterized as a dense focus of cDcp-1 staining adjacent to an array of triangular phalloidin staining of F-actin cones, which together shows a snapshot of the individualization complex removing cytoplasmic components (Fig. 3A’; yellow arrow). Finally, waste bags were characterized as a rotund structure near the apical tip of the testes with co-localized cDcp-1 and phalloidin staining, which is composed of the individualization complex and cytoplasmic components that pinched off after finishing the remodeling of the spermatid^24^ (Fig. 3A’; white arrow). Mutant testes stained with anti-cDcp-1 showed the presence of active executioner caspase, indicating that there was no defects in caspase activation. However, the staining revealed clear anatomical differences between the wild-type and mutant testes. All mutant testes (Fig. 3B-D’) contained fewer cystic bulges and lacked waste bags compared to the control testes (Fig. 3A, A’). Quantification of Mi/PB, PB/*Df(3L)ED4978*, and PB/*Df(3L)BSC284* testes is shown for three different stages of individualization: assembling individualization complex, cystic bulges, and waste bags (Fig. 4A). CG7414 mutant testes lacked waste bags, but individualization complexes still formed suggesting that CG7414 is required for the proper progression, rather than the assembly, of individualization complexes.

**Figure. 3.**
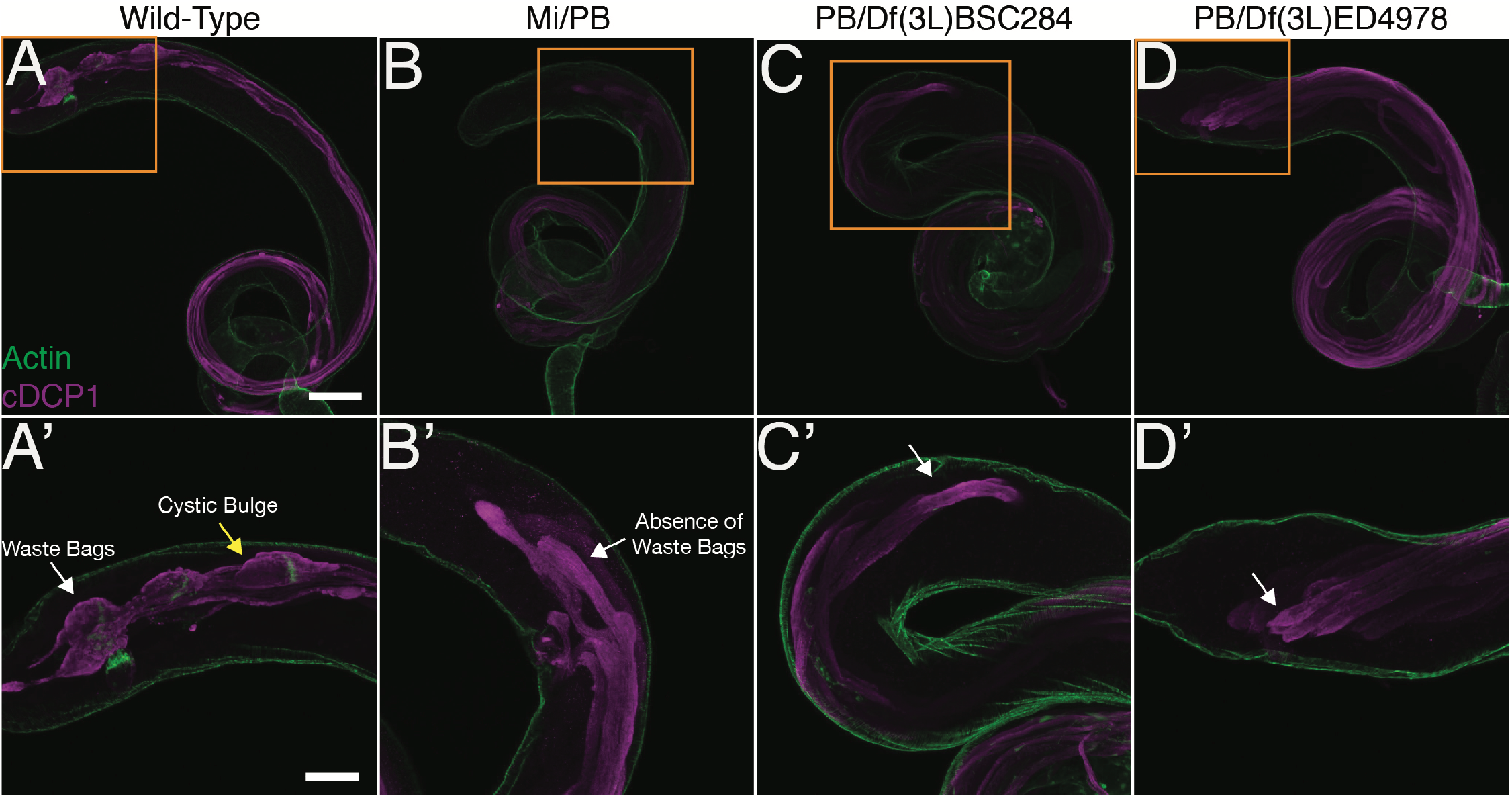
Defective cystic bulges and waste bags in CG7414 mutants. (A) Wild-type testis at 10X magnification (A’) and at 20X magnification. (B) Mi/PB testes are shown at 10X magnification (B’) and at 20X magnification. (C) PB/*Df(3L)BSC284* testes are shown at 10X magnification (C’) and at 20X magnification with a white arrow showing the absence of waste bags. (D) PB/*Df(3L)ED4978* testis are shown at 10X magnification and (D’) at 20X magnification with a white arrow showing the absence of waste bags. (A-D) Orange box outlines the magnified region. Anti-cDcp-1 immunostaining (magenta) and phalloidin-labeled F-actin (green). Scale bars: 100 μM (A-D) and 50 μM (A’-D’).

**Figure. 4.**
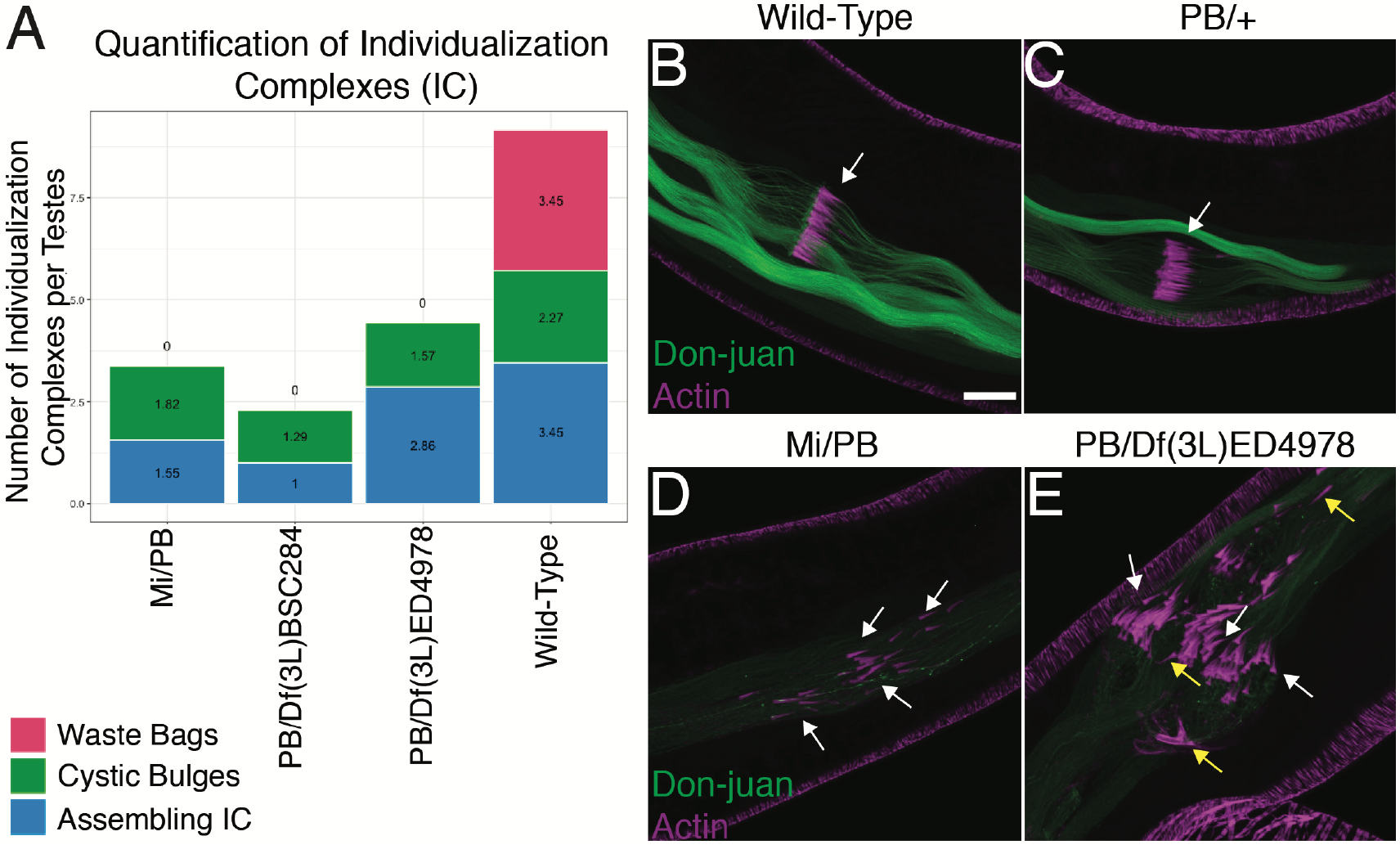
Individualization complex and F-actin cone defects in CG7414 mutants. (A) Quantification of Individualization Complexes of Wild-Type, Mi/PB, PB /Df(3L)BSC284, and PB/Df(3L)ED4978 testes. (B-C) Wild-Type and heterozygous PB/+ testis with a white arrow showing a normal cystic bulge with F-actin cones. (D) Mi/PB testes with white arrows pointing at asynchronous actin cones. (E) PB/Df(3L)ED4978 testis with white arrows pointing at asynchronous F-actin cones and yellow arrows showing reverse polarity of F-actin cones. (B-D) The direction of movement for the F-actin cones is to the left. DJ-GFP is labeled as green and actin staining is labeled as magenta. Images are at 40X magnification. Scale bar: 25 μM.

### CG7414 is required for individualization

F-actin cones drive the individualization complex to remove cytoplasmic contents and to remodel the membranes during separation of individual spermatids^24,30,31^. As expected, WT and heterozygous PB/+ testis showed well-organized, polarized, and aligned F-actin cones that migrate synchronously from the basal to the apical end of the testis. This organized and polarized movement forms the cystic bulges during individualization (Fig. 4 B, C; white arrows). In contrast, both Mi/PB and PB/*Df(3L)ED4978* mutant testes were disorganized. The individualization complexes containing the F-actin cones were not aligned, suggesting a loss of synchrony in their progressive movement (Fig. 4D; white arrows). In addition, some F-actin cones were poorly polarized and occasionally exhibited reversed polarity (Fig 4E; yellow arrows). The abnormal individualization complexes in Mi/PB and PB/*Df(3L)ED4978* mutants ultimately failed to produce waste bags (Fig. 2B-D). These observations suggest that CG7414 mutant testes contained many disorganized spermatids that appeared to be in tangled “knots” (Fig. 4D, E) in contrast to the highly ordered structure of WT spermatids (Fig. 4B, C). We conclude that CG7414, the Drosophila ortholog of EIF2A, is required for individualization.

Although a functional antibody for dEIF2A is not available, we were able to determine the spatial distribution of dEIF5B, the Drosophila ortholog of EIF2A’s interacting partner, during spermatogenesis. In WT testes, we found high levels of eIF5B in the cystic bulges and waste bags consistent with a potential role for local non-AUG translation initiation within these structures^16^ (Fig. 5A, A’). Altogether these observations suggest that CG7414 plays a significant role in Drosophila testes and that non-AUG translation may regulate individualization.

**Figure. 5.**
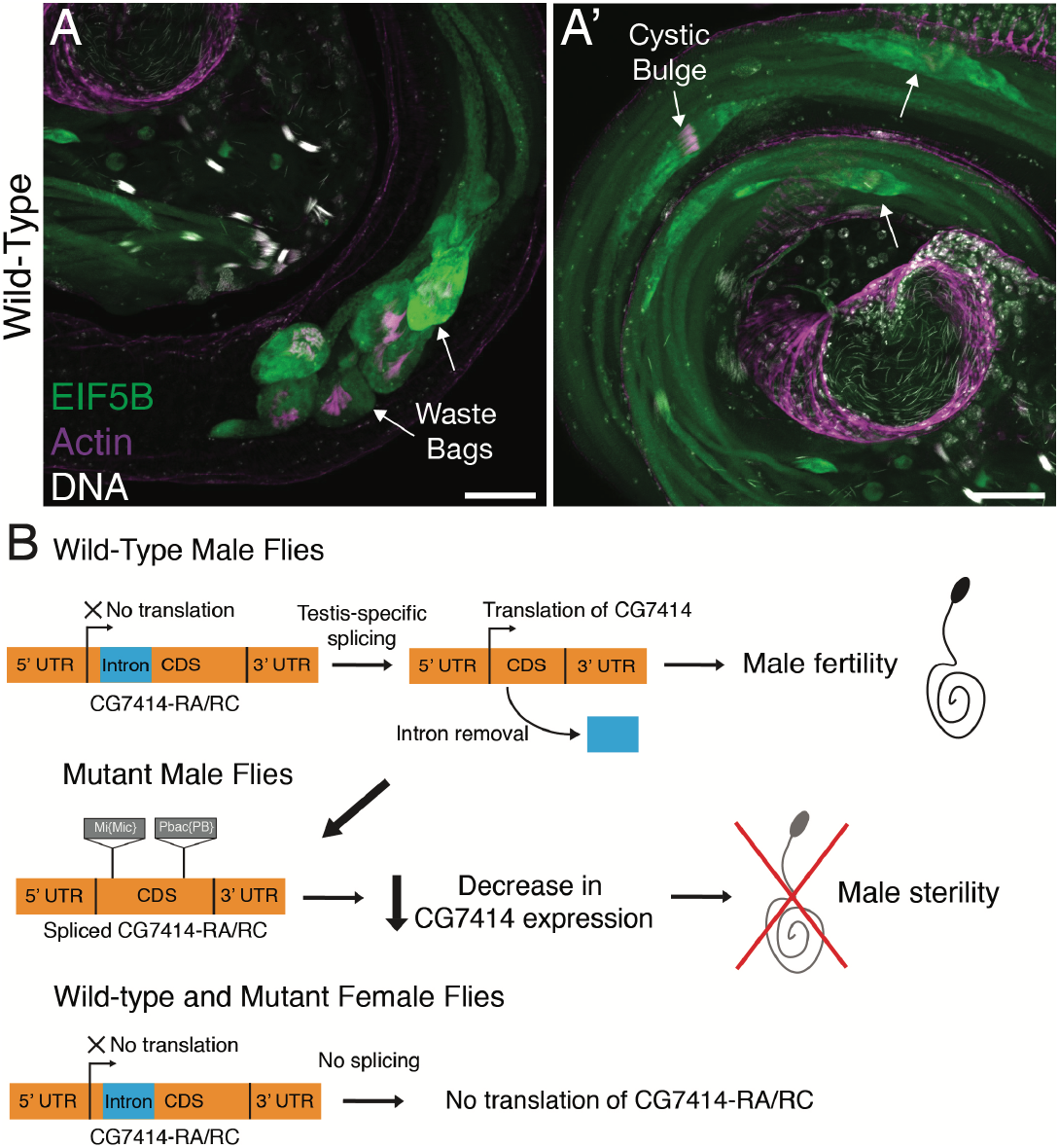
Model for CG7414 function for male fertility. (A) Wild-Type testis shown at 20X magnification with white arrows marking waste bags. (A’) Control testis shown at 20X with white arrows marking cystic bulges. (A – A’) EIF5B is labeled in green, actin is labeled in magenta, and DNA is labeled in white. Scale bar: 50 μM. (B) Wild-type male flies undergo testis-specific splicing for translation of CG7414-RA/RC transcripts. Expression of CG7414 leads to synthesis of proteins needed for individualization. Mutant male flies contain transposon insertions that decrease CG7414 expression leading to male sterility. Wild-type and mutant female flies cannot splice the intron of CG7414-RA/RC leading to no translation.

## Discussion

Fly spermatogenesis has been shown to be a powerful system to uncover the roles of translation initiation factors. In fact, various translation factors have been shown to have testis-specific isoforms with roles in various stages of spermatogenesis including mitosis, meiosis, elongation, and individualization^25,32–33^. However, the roles of unconventional initiation factors, such as the Drosophila ortholog of *EIF2A*, *CG7414*, have not been characterized.

The work reported here demonstrates that dEIF2A is required for normal organism development and viability and that it is essential for male fertility. Furthermore, sex-specific expression of dEIF2A is regulated by sex-specific splicing of the first intron. Consistent with our results, previous work by Wang et al. reported widespread sex-specific intron-retention events^18^. We also found that transcripts encoding both the full-length and truncated forms of dEIF2A were present, the transcript encoding the full-length protein appeared to be more abundant. The truncated protein, if produced at all, would lack the EIF5B binding site. Therefore, it is unclear what function this protein could serve.

Spermatogenesis is an elaborate process that requires numerous unconventional molecular mechanisms including non-lethal caspase activity as well as atypical regulation of protein synthesis^30,31,34^. Our results support a model whereby sex-specific splicing in the testes allows production of PA and/or PC proteins. Together with eIF5B, dEIF2A may initiate local translational from non-canonical CUG or UUG codons, resulting in the synthesis of proteins^16^ that are specifically required to align and polarize the parallel arrays of F-actin cones, which resemble the teeth in a comb. The aligned and polarized cones in turn are necessary for their synchronous migration to separate individual sperm. When the expression of CG7414 is significantly reduced, as in Mi/PB or PB/*Df(3L)BSC284*, we suggest that the failure of local synthesis of spermatogenesis proteins results in the disorganization of the F-actin cones and improper migration of the individualization complexes, leading to spermatid death and thus to male sterility (Fig. 5B). Identification of the relevant translational targets of dEIF2A/eIF5B would shed considerable light on the mechanism.

Asynchronous movement of the actin cones in the CG7414 mutants is consistent with previously characterized phenotypes from cytoskeletal components, such as *jar* (95F myosin) and *mulet* mutants^35–38^, and apoptotic components, such as DRONC and ARK RNAi knockdowns^31^. In addition, post-translational modifying enzymes are integral for the stability of axonemes and individualization, such as the glyclase TTLL3B, which appends glycine onto microtubules^39^.

It is also possible that translation from non-canonical codons is not specifically concentrated in the individualization complex but rather that the apparent accumulation of eIF5B there is a consequence of the concentration of cytoplasmic components. In this case structural integrity of the individualization complex may require components that are translationally regulated in a dEIF2A-dependent manner. Again, identification of the relevant target(s) will shed significant light on the mechanism

Our work thus adds a new insight into the biological functions of non-canonical mRNA translation and into the specific and highly regulated process of spermatogenesis. Spermatogenesis is highly conserved with many similarities between insect and mammalian processes, especially in the utilization of spatiotemporally regulated translation. Further study of the mechanisms of action of this highly conserved protein in Drosophila will likely provide key and general insights into its functions in metazoan animal development, physiology, and reproduction.

## Materials and Methods

### Fly lines and genetics

All flies were kept at 25℃ unless noted separately. w^1118^ were used as Wild-Type or WT. Stocks obtained from the Bloomington Drosophila Stock Center (NIH P40OD018537) were used in this study. The following lines were received from Bloomington: CG7414^c00723^ (BL#10233), CG7414^MI06462^ (BL#42152), Df(3L)BSC284 (BL#23669), Df(3L)ED4978 (BL#8101), EIF5B-GFP (BL#64446), and DJ-GFP (BL#5417). CG7414^c00723^ is labeled as PB since it contains a PiggyBac transposon inserted in the 2nd intron of CG7414. CG7414^MI06462^ is labeled as Mi since it contains a Minos transposon inserted in the 2nd exon of CG7414. EIF5B-GFP contains a transgenic MiMIC based protein cassette which is inserted into the intron of EIF5B for internal protein tagging. DJ-GFP contains a transgenic P-element construct containing a don-juan coding sequence fused to GFP under the endogenous don-juan regulatory sequences inserted in the 2nd chromosome.

### Male fertility assay

7-10 male flies from the desired genotype were isolated after 1-3 days from eclosion. The male fertility assay was performed by mating a single male fly with 2-3 virgin female w^1118^ flies for 10 days, then the flies were removed. Each vial was checked every day for larval progeny and newly eclosed flies, which were counted and sexed. The number of independent crosses or replicates for desired male flies was designated as N. % of fertility was calculated as total vials with progeny / total vials assayed.

### RNA isolation and RT-qPCR

1-3 days old flies were separated by sex and flash frozen in a −80℃ freezer. Frozen flies were incubated with 1 mL of TRIzol Reagent (Life Technologies) and homogenized with a dounce. The homogenized sample was incubated at room temperature for 5 minutes then centrifuged at 12,000x rcf at 4℃ to pellet the exoskeletons and insoluble fractions. The supernatant was removed for TRIzol-Chloroform extraction to isolate total RNA. For reverse transcription, two micrograms of total RNA and 50 picomol of reverse transcription Oligo(dT)_20_ was used to generate first-strand cDNA of polyadenylated mRNAs using the Superscript III First-Strand Synthesis System (Invitrogen). Total RNA was melted with dNTPs and reverse transcription oligonucleotide at 65 °C for 5 min and immediately chilled on ice before proceeding with remaining cDNA synthesis steps. All cDNA samples were treated with RNAse H prior to PCR. One μl of the total cDNA was used for the PCR (20 μl volume) performed with Taq DNA polymerase (New England BioLabs) and primers (Table 2). RP49 or RPL39 is a ribosomal protein that serves as a control for all PCR and qPCR analysis. PCR reactions were run on an agarose gel. For Sanger sequencing, PCR products of interest were excised from gel and gel extracted using the Qiagen Gel Extraction Kit (Qiagen). Sequencing results were aligned to a reference genome utilizing the Benchling cloud software (https://benchling.com). Reverse transcription with quantitative PCR (RT-qPCR) was performed using 1 μl of cDNA as a template with qPCR primers (Table 2) and iTaq Universal SYBR Green Supermix (Biorad) according to the manufacturer’s instruction and analyzed using CFX Connect Real Time System (BioRad). The experimental replicate was measured in technical triplicates. The gene expression of CG7414 was normalized to RP49. The qPCR data was analyzed using the ΔΔCt method to calculate the relative expression of CG7414 transcripts. The absence of the reverse transcriptase or RT(-) with RNAse H treatment served as negative controls for the qPCR assay and RT-PCR. qPCR results were visualized as a bar graph using GraphPad Prism Version 9.0.2 for MacOS, GraphPad Software, San Diego, California USA, www.graphpad.com. Four biological replicates (N=4) were done for each qPCR sample. 2-way ANOVA was performed with the samples grouped by sex.

**Table 2.**
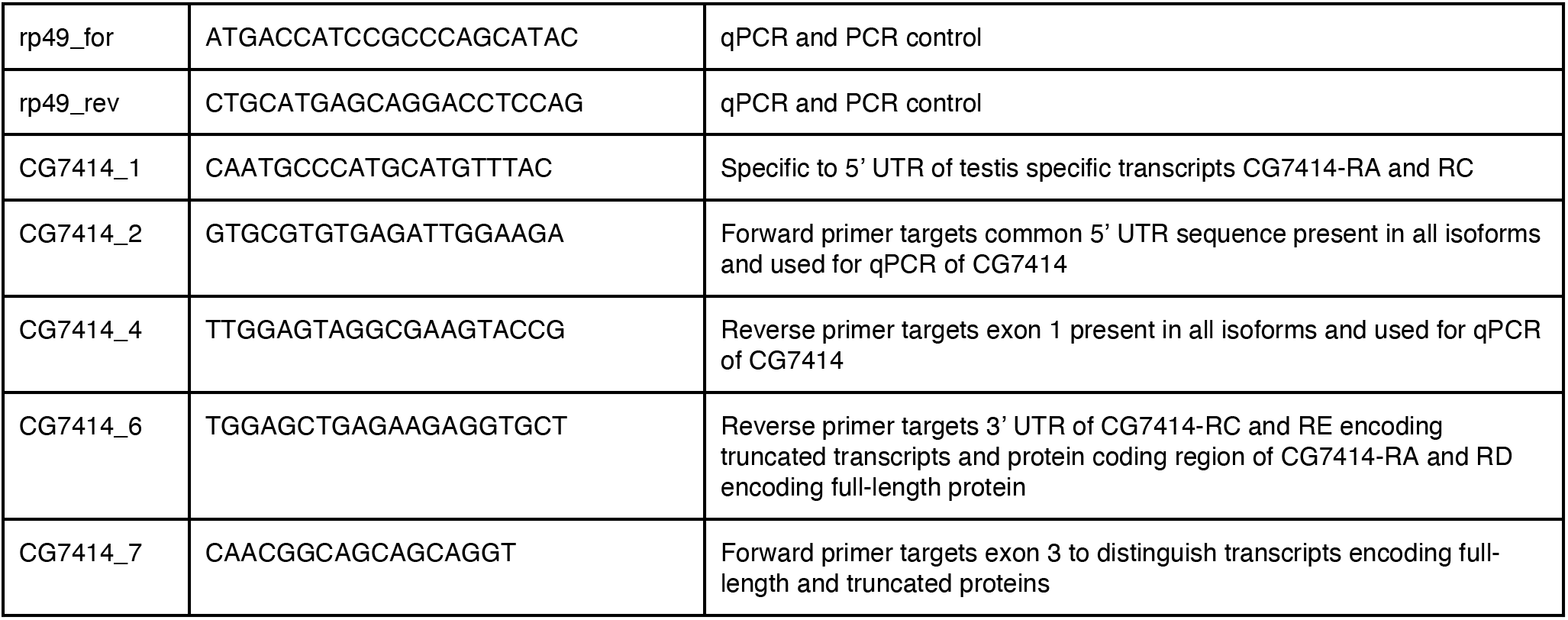
List of primers

### Microscopy

Images were taken on a Zeiss Laser Scanning Microscope 800, using a 10X or 20X magnification 1.2 N.A. objective or 40X magnification 1.4 N.A. water objective. Images were taken at 16 bits with an averaging of 4 on the Zeiss Blue parameter settings. All comparative images were taken with the same laser settings. Fiji was utilized for image analysis and processing.

### Immunohistochemistry

Young adult male flies were collected 1-3 days after eclosion. The testes were dissected in S2R+ medium (supplemented with 10% FBS + 1% Pen-Strep) or 1X PBS and fixed in 4% paraformaldehyde for 20-22 minutes at room temperature. Then, the testes were briefly washed in PBX (1X PBS+0.1% Tween-X100) three times with the final wash for 30 minutes. The testes were blocked in a blocking buffer (PBX + 2% normal goat serum + 3% BSA + 0.02% sodium azide) for 1 hour at room temperature. Samples were incubated with primary antibodies overnight at 4°C. After overnight incubation, the samples were washed in PBX three times and incubated for 2-4 hours with phalloidin to stain F-actin, Hoechst to stain DNA, and fluorescent secondary antibodies, all diluted in blocking buffer. Finally, samples were incubated in Antifade Mounting Medium (Vectashield) overnight at 4°. After overnight incubation, the samples were mounted on a slide for imaging. Primary antibodies include anti-cDcp-1 (1:250 dilution). Secondary antibody conjugated to the fluorophore Alexa-488 and Alexa-568 was used at 1:200. Alexa-657 Phalloidin and Alexa-488 Phalloidin were used at 1:500. Hoechst 33342 was used at 1:1000.

### Quantification of Cystic Bulges and Waste Bags

Original images were opened in Fiji (ImageJ) and pseudo-coloring was utilized to aid in visual identification of individualization complexes. Quantifications were separated into three distinct stages: 1) assembling or assembled Individualization Complexes were based on overlapping nuclei and F-actin staining with needle-shape morphology 2) progressing Individualization Complexes (Cystic bulge) were identified as Individualization Complexes with actin cones and cDcp-1 staining that are still migrating, and 3) progressed Individualization Complexes (Waste Bags) were identified as Individualization Complexes that have finished migrating near the apical end of the testis. Four separate testes were quantified per genotype. Bar graph was created utilizing R Studio and ggplot2 library on the quantification data.

## End Matter

### Author Contributions and Notes

D.D.L and D.J.M designed research, D.D.L. performed research and analyzed data; and D.D.L and D.J.M. wrote the paper. The authors declare no conflict of interest.

## Acknowledgments

The authors would like to thank Izel Tekin, and Dhanajay Thakur for support and assistance with Drosophila work; Jieyan Chen for helpful insights on spermatogenesis; Gongping Sun for the cleaved Dcp-1 antibody; Joe Campanale for imaging assistance and training; members of the Montell lab for support and advice; and C. Ting Wu Lab for material support. D.D.L. was supported by the Genetics and Genomics Training Grant, Harvard University, National Institutes of Health (NIH 5T32GM096911-10) and Undergraduate Research and Creativity Grant, University of California Santa Barbara (#3816, #3401).

---

**Supplementary Figure 1.**
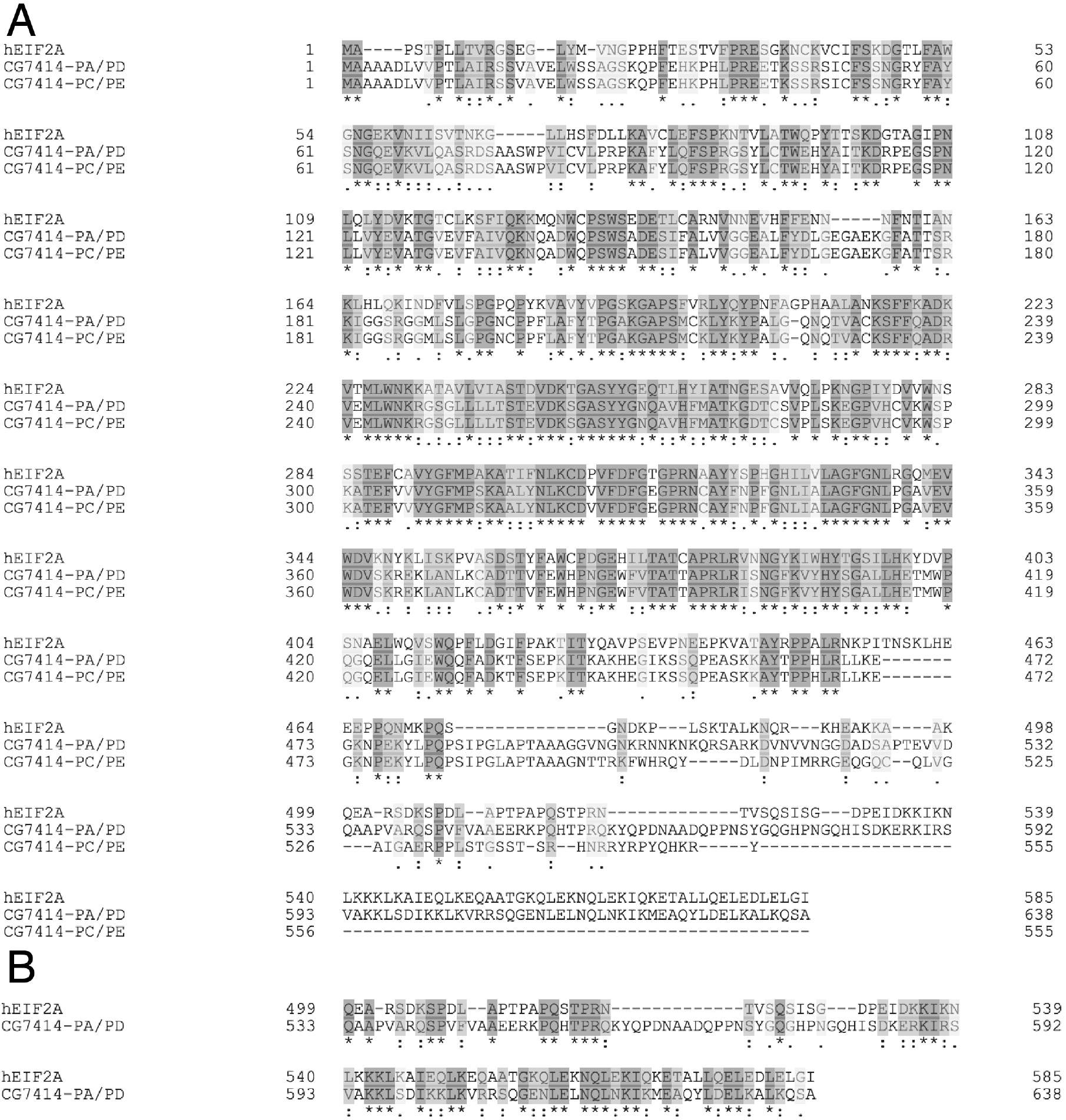
(A) Clustal Omega alignment of the human EIF2A (Q9BY44) with both isoforms of CG7414 (Q9VNX8 and M9MS45). The conserved tRNA binding domain is between residues 232-422 in CG7414 or 216-406 residues in human EIF2A. The EIF5B binding domain is predicted to be located between residues 473-539 in CG7414-PA/PD and residues 473-529 in CG7414-PC/PE, while the known EIF5B binding domain is located between residues 462-503 in human EIF2A. (B) Shows the alignment of human EIF2A (Q9B7FF) and CG7414-PA/PD (QVNX8) for the conserved C-terminus. The conserved mRNA binding domain is located at the C-terminus between residues 545-639 in CG7414-PA/PD and between residues 502-585 in human EIF2A. (A-B) Similar amino acid residues are shown in light gray and identical amino acid residues are shown in dark gray. Fully conserved residues are labeled with an asterisk (*), strongly similar residues are labeled with a colon (:), and weakly similar residues are labeled with a period (.).

**Supplementary Figure 2.**
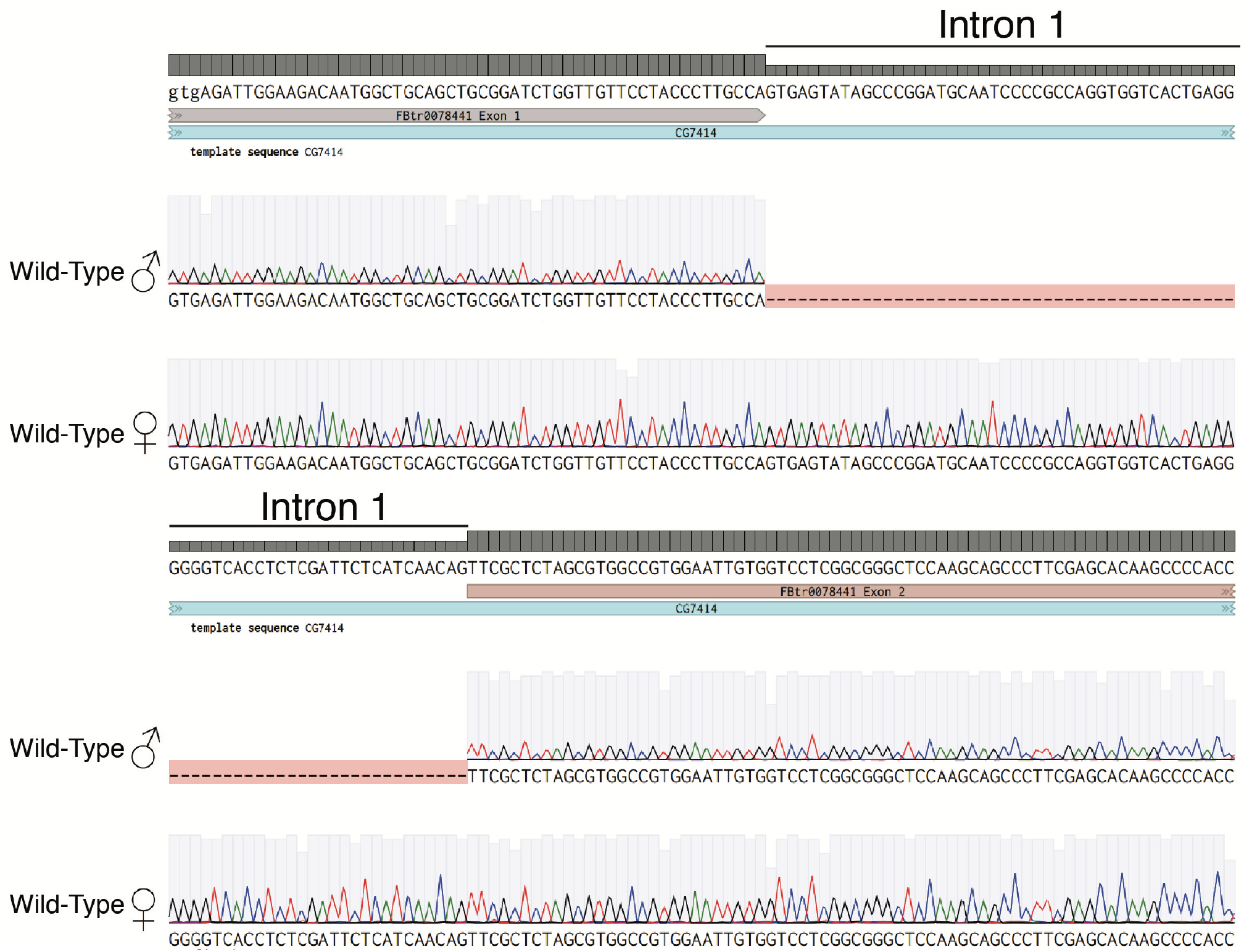
Sanger sequencing of gel extracted PCR products from Lane 2 and 3 of primer pair 1/4 shown in Figure 1D. Shown are the exon-intron junctions for exon 1-intron 1 (above) and intron 1-exon 2 of CG7414 (below). Wild-Type males have no intron sequences, while the females retain intron 1.

**Supplementary Figure 3.**
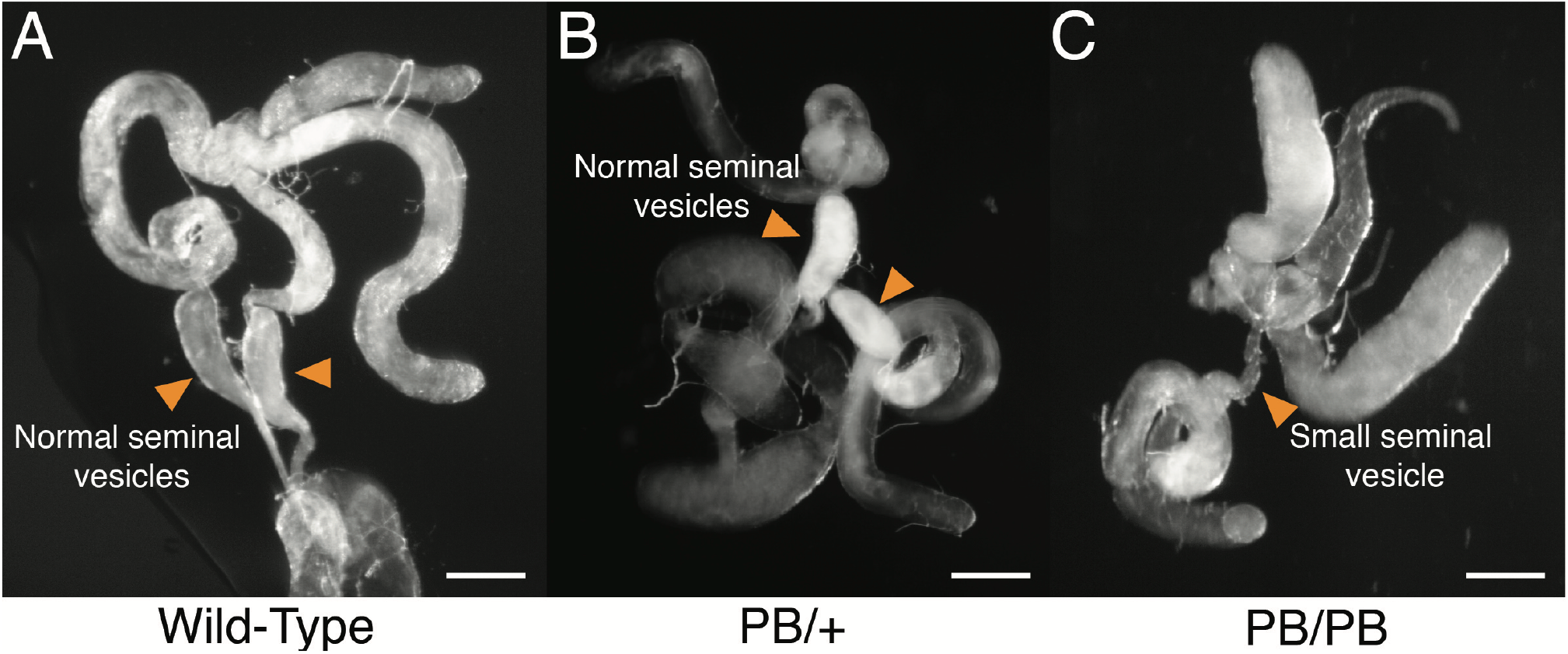
Light microscopy images of (A) 5-day old Wild-Type testes, (B) 5-day old heterozygous PB/+ testes, and (C) 10-day old homozygous PB/PB testes. (A-C) Orange arrows point at seminal vesicles. WT and PB/+ contain large, normal seminal vesicles full of sperm, while the PB/PB testes contain small seminal vesicles with no sperm. Scale bar: 200 μM

## References

1. Sonenberg N, Hinnebusch AG. Regulation of translation initiation in eukaryotes: mechanisms and biological targets. Cell. 2009;136(4):731–745. doi:10.1016/j.cell.2009.01.042

2. Wan J, Qian S-B. TISdb: a database for alternative translation initiation in mammalian cells. Nucleic Acids Res. 2014;42(Database issue):D845–50. doi:10.1093/nar/gkt1085

3. Na CH, Barbhuiya MA, Kim M-S, et al. Discovery of noncanonical translation initiation sites through mass spectrometric analysis of protein N termini. Genome Res. 2018;28(1):25–36. doi:10.1101/gr.226050.117

4. Kearse MG, Wilusz JE. Non-AUG translation: a new start for protein synthesis in eukaryotes. Genes Dev. 2017;31(17):1717–1731. doi:10.1101/gad.305250.117

5. Starck SR, Jiang V, Pavon-Eternod M, et al. Leucine-tRNA initiates at CUG start codons for protein synthesis and presentation by MHC class I. Science. 2012;336(6089):1719–1723. doi:10.1126/science.1220270

6. Starck SR, Tsai JC, Chen K, et al. Translation from the 5’ untranslated region shapes the integrated stress response. Science. 2016;351(6272):aad3867. doi:10.1126/science.aad3867

7. Sendoel A, Dunn JG, Rodriguez EH, et al. Translation from unconventional 5’ start sites drives tumour initiation. Nature. 2017;541(7638):494–499. doi:10.1038/nature21036

8. Trulley P, Snieckute G, Bekker-Jensen D, et al. Alternative Translation Initiation Generates a Functionally Distinct Isoform of the Stress-Activated Protein Kinase MK2. Cell Rep. 2019;27(10):2859–2870.e6. doi:10.1016/j.celrep.2019.05.024

9. Beerman RW, Jongens TA. A non-canonical start codon in the Drosophila fragile X gene yields two functional isoforms. Neuroscience. 2011;181:48–66. doi:10.1016/j.neuroscience.2011.02.029

10. Lorsch JR, Dever TE. Molecular view of 43 S complex formation and start site selection in eukaryotic translation initiation. J Biol Chem. 2010;285(28):21203–21207. doi:10.1074/jbc.R110.119743

11. Gallie DR. The cap and poly(A) tail function synergistically to regulate mRNA translational efficiency. Genes Dev. 1991;5(11):2108–2116. doi:10.1101/gad.5.11.2108

12. Wells SE, Hillner PE, Vale RD, Sachs AB. Circularization of mRNA by eukaryotic translation initiation factors. Mol Cell. 1998;2(1):135–140. doi:10.1016/s1097-2765(00)80122-7

13. Hinnebusch AG. The scanning mechanism of eukaryotic translation initiation. Annu Rev Biochem. 2014;83:779–812. doi:10.1146/annurev-biochem-060713-035802

14. Jackson RJ, Hellen CUT, Pestova TV. The mechanism of eukaryotic translation initiation and principles of its regulation. Nat Rev Mol Cell Biol. 2010;11(2):113–127. doi:10.1038/nrm2838

15. Pestova TV, Kolupaeva VG, Lomakin IB, et al. Molecular mechanisms of translation initiation in eukaryotes. Proc Natl Acad Sci USA. 2001;98(13):7029–7036. doi:10.1073/pnas.111145798

16. Kim E, Kim JH, Seo K, et al. eIF2A, an initiator tRNA carrier refractory to eIF2α kinases, functions synergistically with eIF5B. Cell Mol Life Sci. 2018;75(23):4287–4300. doi:10.1007/s00018-018-2870-4

17. Leader DP, Krause SA, Pandit A, Davies SA, Dow JAT. FlyAtlas 2: a new version of the Drosophila melanogaster expression atlas with RNA-Seq, miRNA-Seq and sex-specific data. Nucleic Acids Res. 2018;46(D1):D809–D815. doi:10.1093/nar/gkx976

18. Wang M, Branco AT, Lemos B. The Y Chromosome Modulates Splicing and Sex-Biased Intron Retention Rates in Drosophila. Genetics. 2018;208(3):1057–1067. doi:10.1534/genetics.117.300637

19. Horowitz H, Berg CA. Aberrant splicing and transcription termination caused by P element insertion into the intron of a Drosophila gene. Genetics. 1995;139(1):327–335.

20. White-Cooper H, Schäfer MA, Alphey LS, Fuller MT. Transcriptional and post-transcriptional control mechanisms coordinate the onset of spermatid differentiation with meiosis I in Drosophila. Development. 1998;125(1):125–134.

21. Blümer N, Schreiter K, Hempel L, et al. A new translational repression element and unusual transcriptional control regulate expression of don juan during Drosophila spermatogenesis. Mech Dev. 2002;110(1-2):97–112.

22. Schäfer M, Nayernia K, Engel W, Schäfer U. Translational control in spermatogenesis. Dev Biol. 1995;172(2):344–352. doi:10.1006/dbio.1995.8049

23. Tokuyasu KT. Dynamics of spermiogenesis in Drosophila melanogaster. VI. Significance of “onion” nebenkern formation. J Ultrastruct Res. 1975;53(1):93–112. doi:10.1016/s0022-5320(75)80089-x

24. Arama E, Agapite J, Steller H. Caspase activity and a specific cytochrome C are required for sperm differentiation in Drosophila. Dev Cell. 2003;4(5):687–697. doi:10.1016/s1534-5807(03)00120-5

25. Ghosh S, Lasko P. Loss-of-function analysis reveals distinct requirements of the translation initiation factors eIF4E, eIF4E-3, eIF4G and eIF4G2 in Drosophila spermatogenesis. PLoS One. 2015;10(4):e0122519. doi:10.1371/journal.pone.0122519

26. Hempel LU, Rathke C, Raja SJ, Renkawitz-Pohl R. In Drosophila, don juan and don juan like encode proteins of the spermatid nucleus and the flagellum and both are regulated at the transcriptional level by the TAF II80 cannonball while translational repression is achieved by distinct elements. Dev Dyn. 2006;235(4):1053–1064. doi:10.1002/dvdy.20698

27. Noguchi T, Lenartowska M, Rogat AD, Frank DJ, Miller KG. Proper cellular reorganization during Drosophila spermatid individualization depends on actin structures composed of two domains, bundles and meshwork, that are differentially regulated and have different functions. Mol Biol Cell. 2008;19(6):2363–2372. doi:10.1091/mbc.e07-08-0840

28. Kaplan Y, Gibbs-Bar L, Kalifa Y, Feinstein-Rotkopf Y, Arama E. Gradients of a ubiquitin E3 ligase inhibitor and a caspase inhibitor determine differentiation or death in spermatids. Dev Cell. 2010;19(1):160–173. doi:10.1016/j.devcel.2010.06.009

29. Noguchi T, Miller KG. A role for actin dynamics in individualization during spermatogenesis in Drosophila melanogaster. Development. 2003;130(9):1805–1816. doi:10.1242/dev.00406

30. Zhong L, Belote JM. The testis-specific proteasome subunit Prosalpha6T of D. melanogaster is required for individualization and nuclear maturation during spermatogenesis. Development. 2007;134(19):3517–3525. doi:10.1242/dev.004770

31. Huh JR, Vernooy SY, Yu H, et al. Multiple apoptotic caspase cascades are required in nonapoptotic roles for Drosophila spermatid individualization. PLoS Biol. 2004;2(1):E15. doi:10.1371/journal.pbio.0020015

32. Baker CC, Fuller MT. Translational control of meiotic cell cycle progression and spermatid differentiation in male germ cells by a novel eIF4G homolog. Development. 2007;134(15):2863–2869. doi:10.1242/dev.003764

33. Marygold SJ, Attrill H, Lasko P. The translation factors of Drosophila melanogaster. Fly (Austin). 2017;11(1):65–74. doi:10.1080/19336934.2016.1220464

34. Bader M, Arama E, Steller H. A novel F-box protein is required for caspase activation during cellular remodeling in Drosophila. Development. 2010;137(10):1679–1688. doi:10.1242/dev.050088

35. Fabrizio JJ, Hime G, Lemmon SK, Bazinet C. Genetic dissection of sperm individualization in Drosophila melanogaster. Development. 1998;125(10):1833–1843.

36. Fabrizio JJ, Aqeel N, Cote J, et al. Mulet (mlt) encodes a tubulin-binding cofactor E-like homolog required for spermatid individualization in Drosophila melanogaster. Fly (Austin). 2012;6(4):261–272. doi:10.4161/fly.21533

37. Nuwal T, Kropp M, Wegener S, Racic S, Montalban I, Buchner E. The Drosophila homologue of tubulin-specific chaperone E-like protein is required for synchronous sperm individualization and normal male fertility. J Neurogenet. 2012;26(3-4):374–381. doi:10.3109/01677063.2012.731119

38. Hicks JL, Deng WM, Rogat AD, Miller KG, Bownes M. Class VI unconventional myosin is required for spermatogenesis in Drosophila. Mol Biol Cell. 1999;10(12):4341–4353. doi:10.1091/mbc.10.12.4341

39. Rogowski K, Juge F, van Dijk J, et al. Evolutionary divergence of enzymatic mechanisms for posttranslational polyglycylation. Cell. 2009;137(6):1076–1087. doi:10.1016/j.cell.2009.05.020

